# Inhibition of MreB and ftsZ proteins to minimize *E. coli* biofilms formation

**DOI:** 10.1101/523167

**Authors:** Emile Charles

## Abstract

In the United States, more than two million individuals become infected by antibiotic-resistant bacteria, resulting in over 23,000 deaths annually. Bacterial biofilms, one of the major causes of this resistance, form a complex extracellular matrix that physically block antibiotic treatment. Within planktonic bacteria, two proteins, MreB and ftsZ, play a key role in bacterial cell growth and development. MreB regulates this development through maintaining the rod-like shape of gram-negative bacteria, while ftsZ regulates the timing and location of cell division. The present study compared the effects of two protein-inhibitors on biofilm formation of *E. coli*; the inhibitors, A22 Hydrochloride and PC190723, inhibit MreB (cell shape) and ftsZ (cell division), respectively. Efficacy was measured with a crystal violet staining assay. Four experiments were designed testing 1) the minimum inhibitory concentration of the inhibitors, 2) the synergistic effect of the inhibitors, 3) the microscopic effects of the inhibitors, and 4) the effect of the inhibitors on antibiotic susceptibility. A mid-level dosage of A22 significantly decreased biofilm density while there was no response to PC190732. The effect of A22 was verified microscopically, observing the change from bacilli cells to coccoid ones via the inhibition of MreB. In the second experiment, with conjunct inhibition, no interaction was found. Lastly, A22 was as effective as Amoxicillin in disrupting biofilms. The inhibition of MreB was found to have a key role in biofilm development. A model is proposed for biofilm density based on cell shape as affected by MreB.

**Importance:** Each year, more than 2 million Americans acquire antibiotic-resistant infection and 23,000 of them die (CDC, 2013). In a study done by Barsoumian et. al (2015), there was a 16% mortality rate pertaining to biofilm-related infections while non-biofilm infection caused a 5% mortality rate. These casualties aren’t limited to the United States. Abroad, antibiotic resistance is a huge issue: 25,000 deaths estimated in the EU; 38,000 deaths in Thailand; and 58,000 deaths in India, among infants alone (CDC, 2012). It is these statistics that inform us that antibiotic resistance must be addressed.

## Introduction

Antibiotic resistance is a global pandemic. One cause of this resistance is the formation of biofilms. Biofilms, a collection of bacterial cells embedded in an extracellular matrix, adhere to surfaces. The biofilm matrix act as a physical means of antibiotic tolerance in that antibiotics are unable to penetrate beyond the surface layer of cells (Kumar et al., 2017). Biofilms cause antibiotics, and other medicinal treatment, to be practically ineffective (Costerton et al., 1999). Biofilms, once formed, cause the bacterial cells to become encapsulated, which leads to the difficulty in treatment. One of the largest causes of nosocomial infections is *Pseudomonas aeruginosa*, a common but facultative harmful bacteria that can cause multiple organ system failure and mortality once it has formed a biofilm (Rasamiravaka et al., 2015). *P. aeruginosa* (Hoyle et al., 1992) produced biofilms considered to be impenetrable by antibiotics (Anderl et al., 2000). The issue of biofilm formation and, subsequently, antibiotic resistance, presents a problem. Methods to break apart, change the structure of, and damage the biofilms are currently being researched. The issue is no longer what antibiotic can best affect bacteria, but rather what can most effectively get inside the biofilm and can prevent biofilm formation in the first place.

This study examined two proteins that play a key role in the development of biofilms. One of these proteins is MreB, acting as an elongation factor within gram-negative bacteria (Bonez et al., 2017). When MreB is present, bacterial cells are shown to have a rod-like, or bacillus, structure. In the absence of this protein, cells are shown to change to more circular shape (Bonez et al., 2017). This shape-change, from a bacillus shape to that of a coccoid one, is caused by a depolymerization of MreB—a structural protein that would otherwise support the elongated shape of the cells (Bonez et al., 2017). The second protein is ftsZ, which may also affect biofilm formation. FtsZ plays a significant role in the process of cell division. The protein first acts when the septum of division is forming. As the cell elongates, the protein spreads in a ring, commonly known as the “Z-ring”, around the middle of the cell (Margolin, 2005). After the ring has formed, multiple other proteins are recruited to help assist with the final steps of cellular division, ending with cytokinesis (Margolin, 2005).

One strategy to counter biofilm growth is the inhibition of theses morphology-determining proteins. A22 Hydrochloride is a compound found to have an inhibitory effect on MreB (Bonez et al., 2017). When A22 is applied, cells are shown to undergo the previously described morphological change, from bacillus to coccoid (Bonez et al., 2017). The shape change may allow for biofilms to become more porous and more susceptible to treatment. PC190723, a benzoic acid derivative, is also shown to have an inhibitory effect (Andreu et al., 2010). Acting on ftsZ, PC190723 effectively inhibits the protein and causes for the division cycle to be altered (Brown et al., 2008). The inhibitor works to denature ftsZ from a multi-stranded protein to a single-stranded one, weakening and disrupting cell division (Andreu et al., 2010).

Manipulating biofilm formation and structure have great potential for addressing the antibiotic tolerance features of biofilms and can be done through the uses of inhibitory compounds such as A22 and PC190723, which affect cell shape and cell division, respectively. This study investigated the use of the two inhibitory compounds, A22 and PC190723. The two compounds’ inhibitory effect were measured, alone and in combination. The effectiveness was compared to the effectiveness of an antibiotic, as well as the increased susceptibility of cells affected by them. Microscopy was done to examine the morphological effects of the proteins.

## Methods

### Biofilm Growth

A bacterial culture of *Escherichia coli* was grown overnight within 100 mL of Luria Broth in a 250 mL flask. The culture was incubated in a shaking water bath set at 37° C. The following day, the culture was diluted with M9 minimal salts to a 1:100 ratio. The M9 was supplemented with 2% glucose and 1 mM MgSO_4_. 10µL of the culture was added to 1 mL of the M9 within each well of a 24-well plate. Prior to aliquoting the culture sample from the flask, the flask was vortex as to not allow for settling of bacteria. The 24-well plates grew at 37° C for 72 hours to establish the biofilm.

### Concentration Experiment

The first Concentration experiment was used to evaluate the minimum inhibitory concentration (MIC) of the two chemicals on biofilm formation. A control (zero), low, medium, and high dosage of inhibitory chemical was added to the respective wells. Treatments were applied prior to the 72-hour incubation period. Within the plates, each column of six wells was treated with a concentration. The sample size per treatment was 6. The experimental unit was a well. The experimental design of the concentration experiment is shown in Figure 1. The treatment concentrations of the two chemicals are shown in Table 1.

**Table 1.** Dosage treatment of 2 protein inhibitors.

| Concentration | A22 (µg/mL) | PC190723 (µg/mL) |
| --- | --- | --- |
| Control | 0 | 0 |
| Low | 100 | 100 |
| Medium | 200 | 150 |
| High | 400 | 200 |

**Fig 1.**
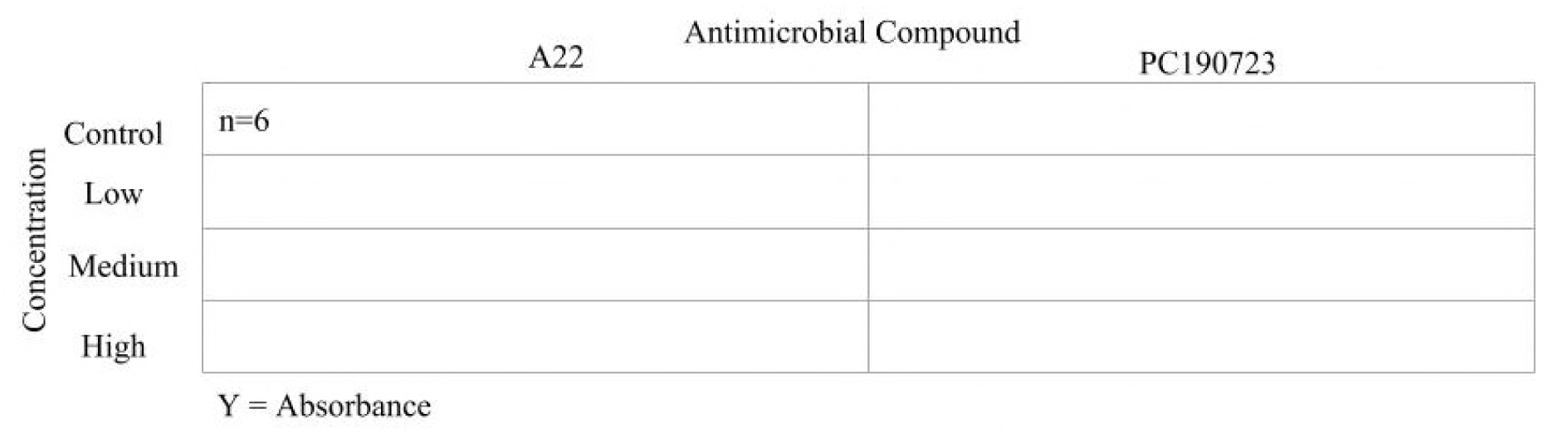
Experimental Design of Concentration Experiment. Treatment, Treatment Level, Sample Size, and Response Variables are shown.

### Combination Experiment

The MIC determined from the previous experiment determined the treatments of the Combination experiment. The Combination experiment was designed to determine whether there were additive or synergistic effects of the two chemicals. The four treatments, a control (zero), A22, PC, and A22+PC, compared the effect of the combination of the two inhibitors to the effect of the two, individually. The treatments were applied prior to the 72-hour incubation period. The sample size per treatment was 18. The experimental unit was a well. The experimental design of the second experiment is shown in Figure 2.

**Fig 2.**
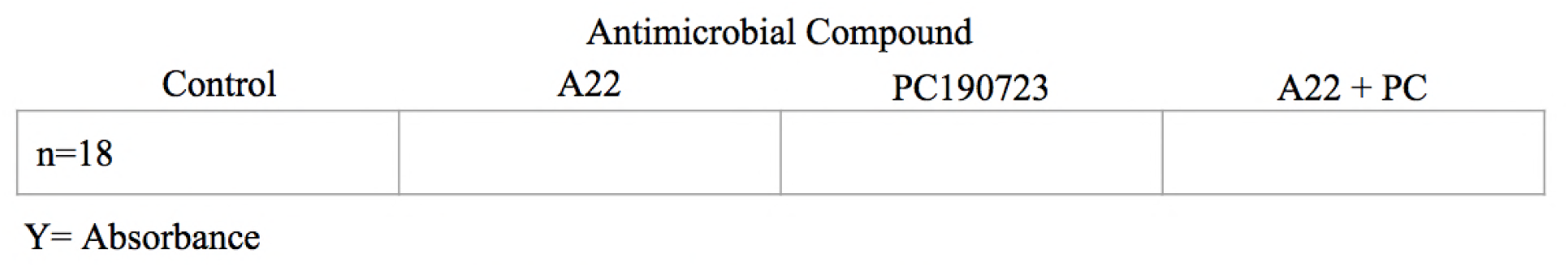
Experimental Design of Combination Experiment. Treatment, Sample Size, and Response Variables are shown.

### Antibiotic Experiment

The third and final experiment was the Antibiotic experiment. The Antibiotic experiment built off the previous two experiments. The treatments included a control (zero), a protein inhibitor (A22), Amoxicillin, and a combination of Amoxicillin and inhibitor. The concentration of the protein inhibitor was the MIC determined from the Concentration experiment. The dosage of Amoxicillin was 2 µg/mL, the accepted MIC of Amoxicillin against *E. coli* (IDEXX Reference Laboratories, 2013). The A22 was applied prior to the 72-hour incubation period. The Amoxicillin was applied at time 48-hours after incubation. The sample size per treatment was 18. The experimental unit was a well. The experimental design of the third experiment is shown in Figure 3.

**Fig 3.**
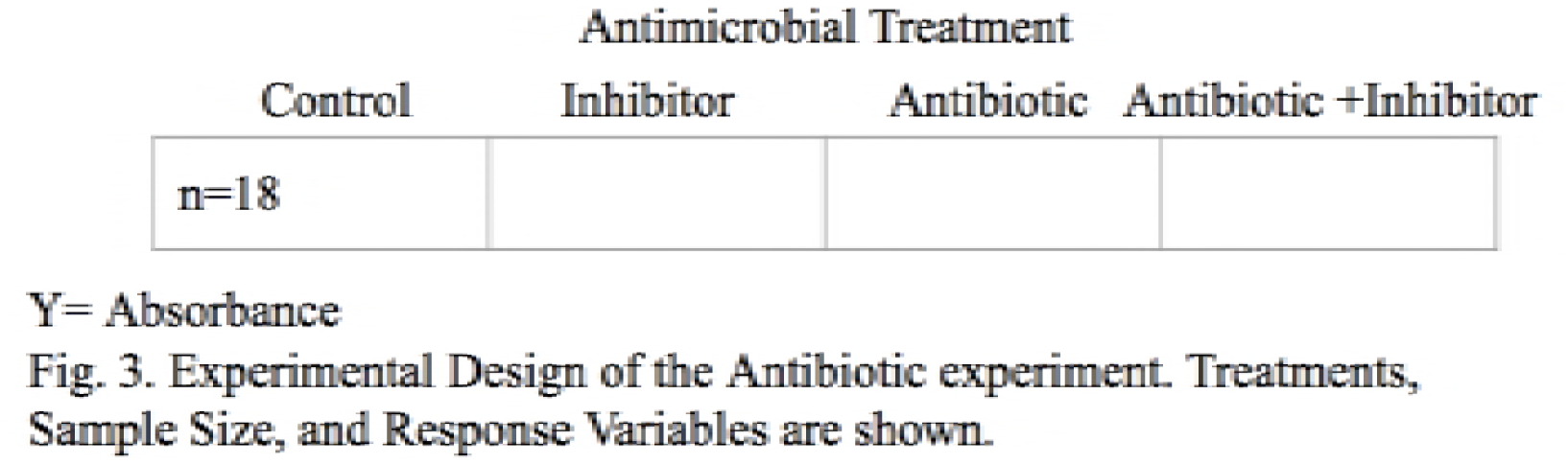
Experimental Design of the Antibiotic experiment. Treatments, Sample Size, and Response Variables are shown.

### Crystal Violet Assay

A crystal violet assay was used to quantify the biofilm growth. Following the 72-hour period of the experiments, the assay called for the well plates to be washed out in lukewarm water to rid the wells of any planktonic bacteria. After approximately 15 minutes of the well-plates air drying, 1 mL of crystal violet (CV) was added to each well. The CV remained in the plates for 15 minutes before it was similarly washed off in lukewarm water. Figure 4 is an image of a biofilm within the well-plate post-staining stage. The CV was solubilized by adding 3 mL of ethanol to each well. Another 15 minutes was given for the stain to detach from the plate. Spectrophotometry was done of the solutions within each plate, quantifying the biofilms within each well through the measure of absorbance. This was done using a Thermo-Scientific Spectronic 200 spectrophotometer set to a wavelength of 560 nanometers. This assay was adapted from that of Agile Sciences, Inc (D. Zeng, 2018).

**Figure 4.**
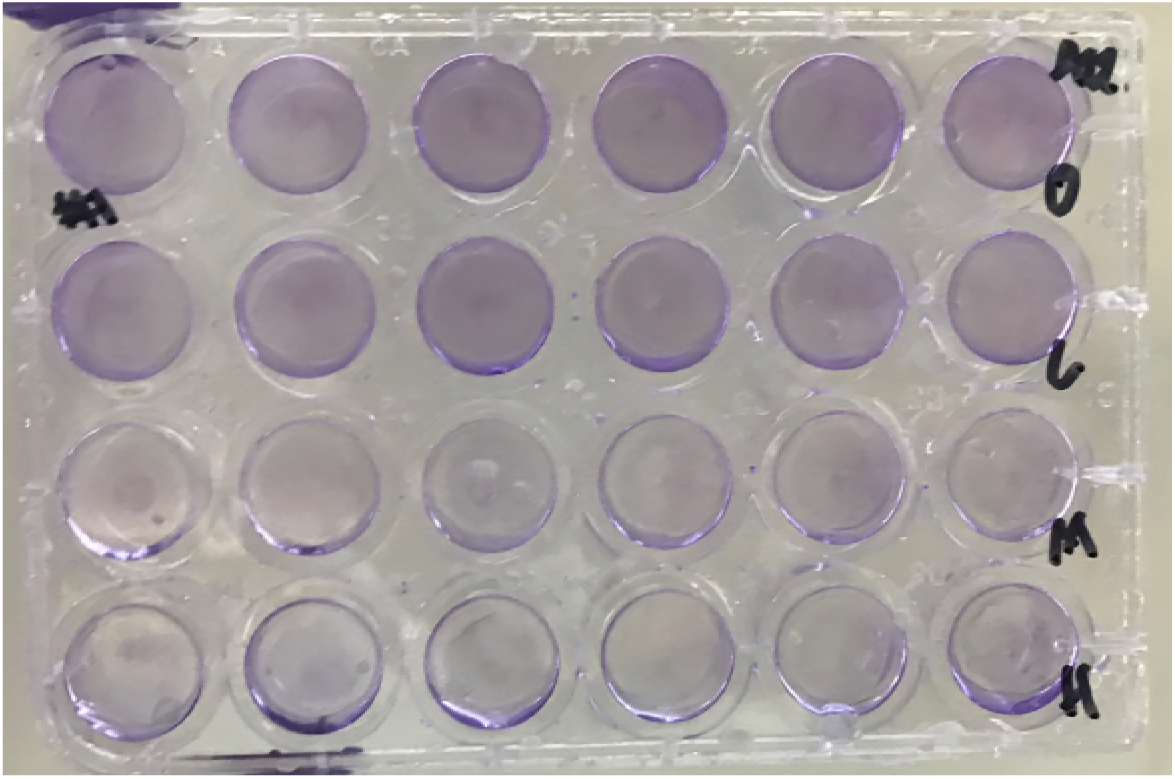
24-well plate stained with crystal violet.

### Microscopy

Microscopy was also done to examine the morphological effects of the A22 and PC190723 compounds. The drugs were applied similarly over the 72 hours period, yet a trypan blue assay was used instead of the CV. An aliquot of about 3 mL from the well plate of normal, A22-treated, and PC190723-treated *E. coli* cells were heat fixed onto separate microscope slides, stained with trypan blue, and viewed under a microscope at 400x. Excess stain was washed off with deionized water, as to not obscure the view. This test was to determine whether the inhibitors affected cell shape.

### Data Analysis

Data were analyzed in JMP 10 with all figured created in Excel. An Analysis of Variance (ANOVA) test was done as well as a Tukey’s mean separation test.

## Results

### Concentration Experiment

Of the A22, it was determined that that was a significant difference between the Control and Medium treatment (ANOVA, df=3, p=.0188). See Table 2. Significant difference was detected between the control and the high treatments, yet no significance was seen between the medium and high treatments (Tukeys). The results of the A22 treatment is shown in figure 5. The MIC of A22 was determined to be the medium dosage (200 µg/mL).

**Table 2.**
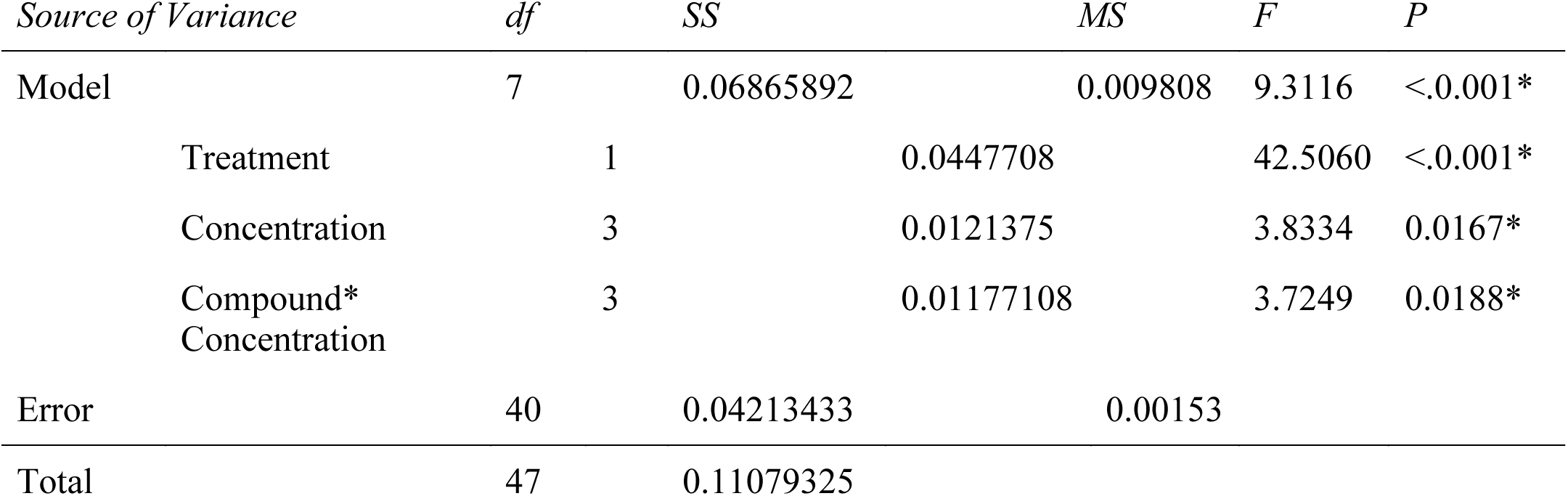
ANOVA results from Concentration Experiment. Asterix (*) represent significant values(p<.05).

**Figure 5.**
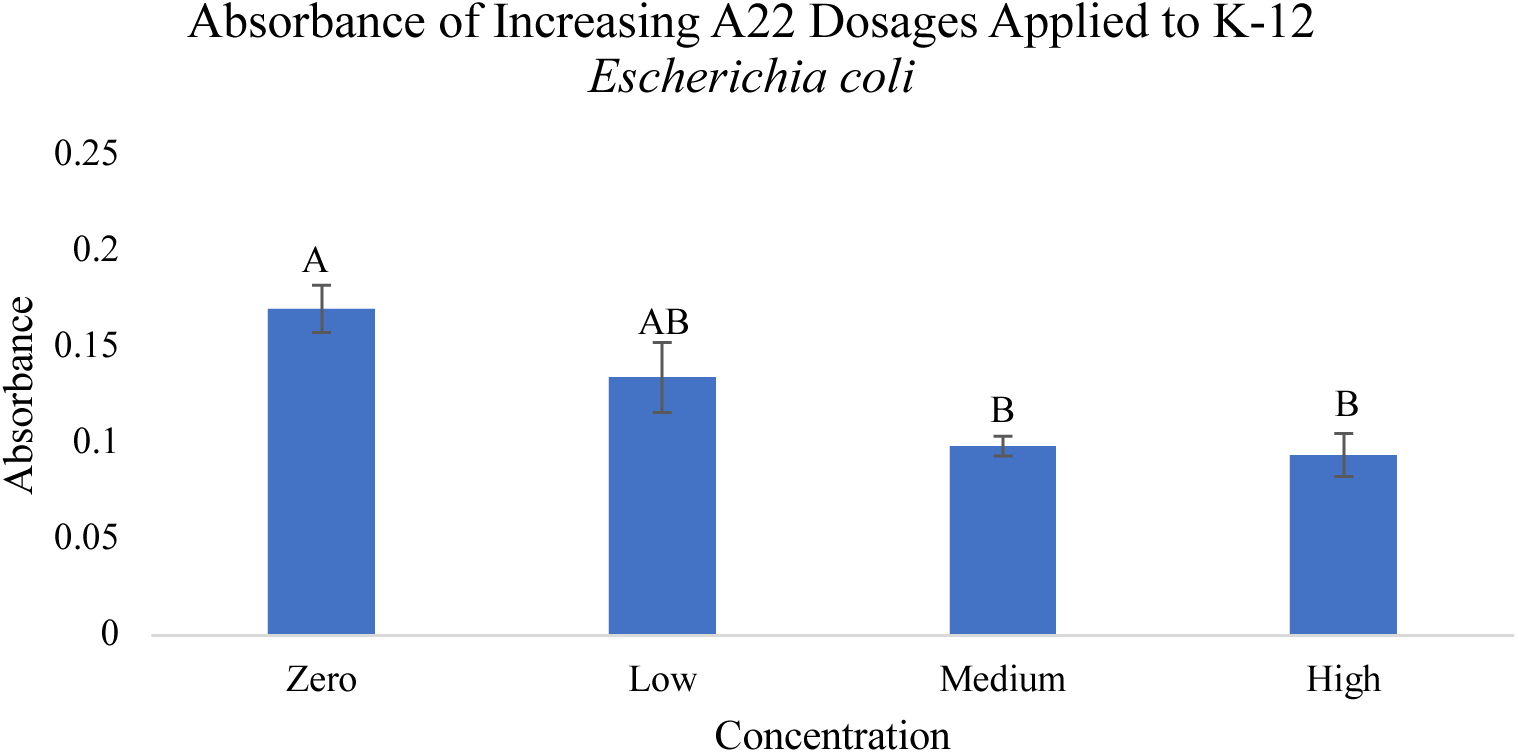
Mean absorbance values of crystal violet solutions within A22 treated well plates. Treatments are the following: 0 µg/mL, 100 µg/mL, 200 µg/mL, and 400 µg/mL. Error bars represent one standard error from the mean. Letters represent Tukey’s mean separation

Within the PC190723 treatment, it was determined that there was no dose response. All four of the dosages were statistically indistinguishable, including the negative control. The low concentration was used as the MIC moving forward in order to differ from the control. The data from this experiment are shown in Figure 6.

**Figure 6.**
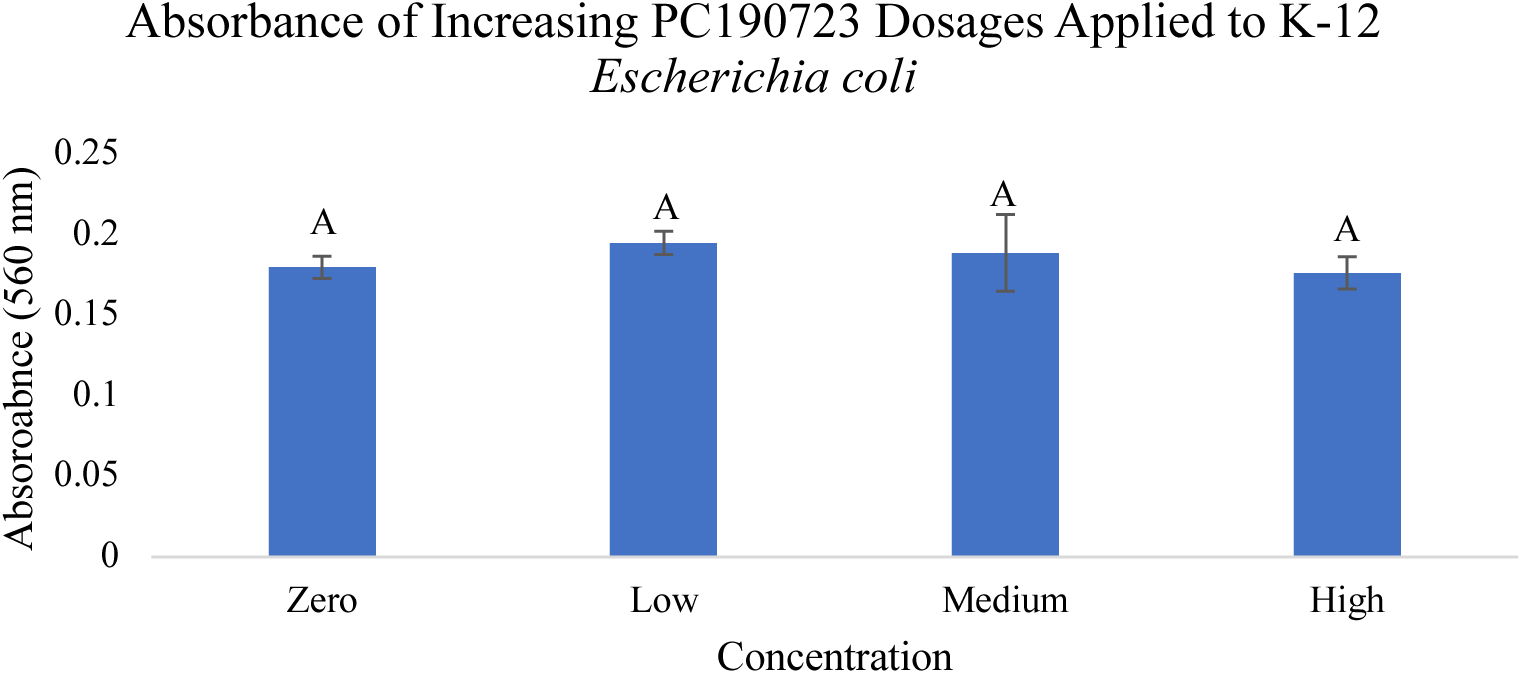
Mean absorbance values of crystal violet solutions within PC190723 treated well plates. Treatments are the following: 0 µg/mL, 100 µg/mL, 150 µg/mL, and 200 µg/mL Error bars represent one standard error from the mean. Letters represent Tukey’s mean separation.

See Table 2 and Figure 7 for complete results.

**Figure 7.**
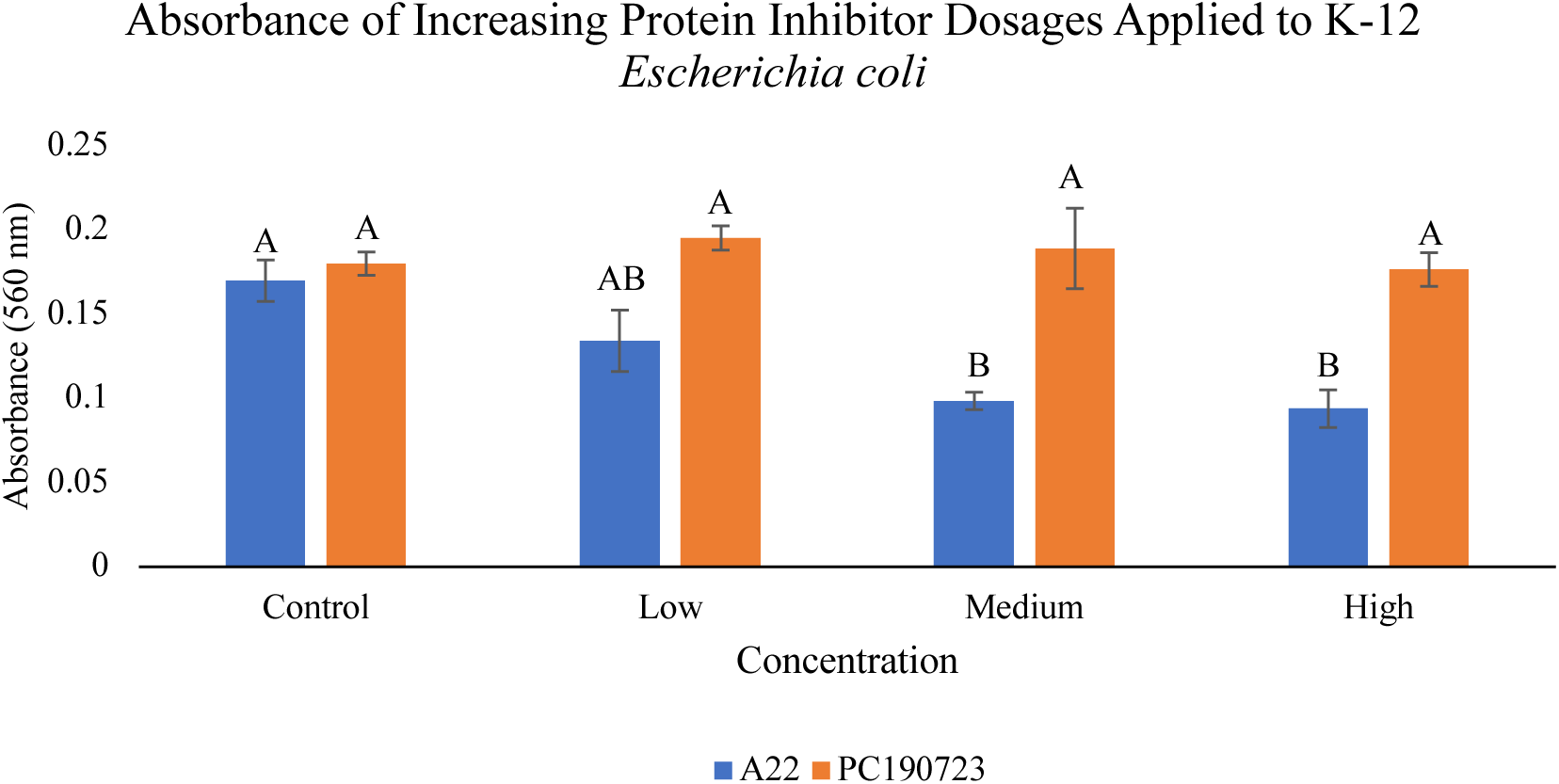
Mean absorbance values of crystal violet solutions within PC190723 treated well-plates. Treatments are the following: Control (Zero), Low, Medium, and High. The specific dosage levels are found in Table 1. Error bars represent one standard error from the mean. Letters represent Tukey’s mean separation.

### Combination Experiment

As expected from the initial experiment, the PC190723 treatment continued to have no effect. Individually, the PC treatment was not statistically different from the control treatment. Additionally, the A22 treatment was not statistically different from the combination treatment. See Figure 8 and Table 3.

**Table 3.** ANOVA results from Combination Experiment. Asterix (*) represent significant values (p<.05).

| *Error* | *dF* | *SS* | *MS* | *F* | *P* |
| --- | --- | --- | --- | --- | --- |
| Model | 3 | 0.13727682 | 0.045759 | 62.8902 | <.0001* |
| Treatment | 3 | 0.13727682 |  | 62.8902 | <.0001* |
| Error | 68 | 0.04947683 | 0.0006728 |  |  |
| Total | 71 | 0.18675365 |  |  |  |

**Figure 8.**
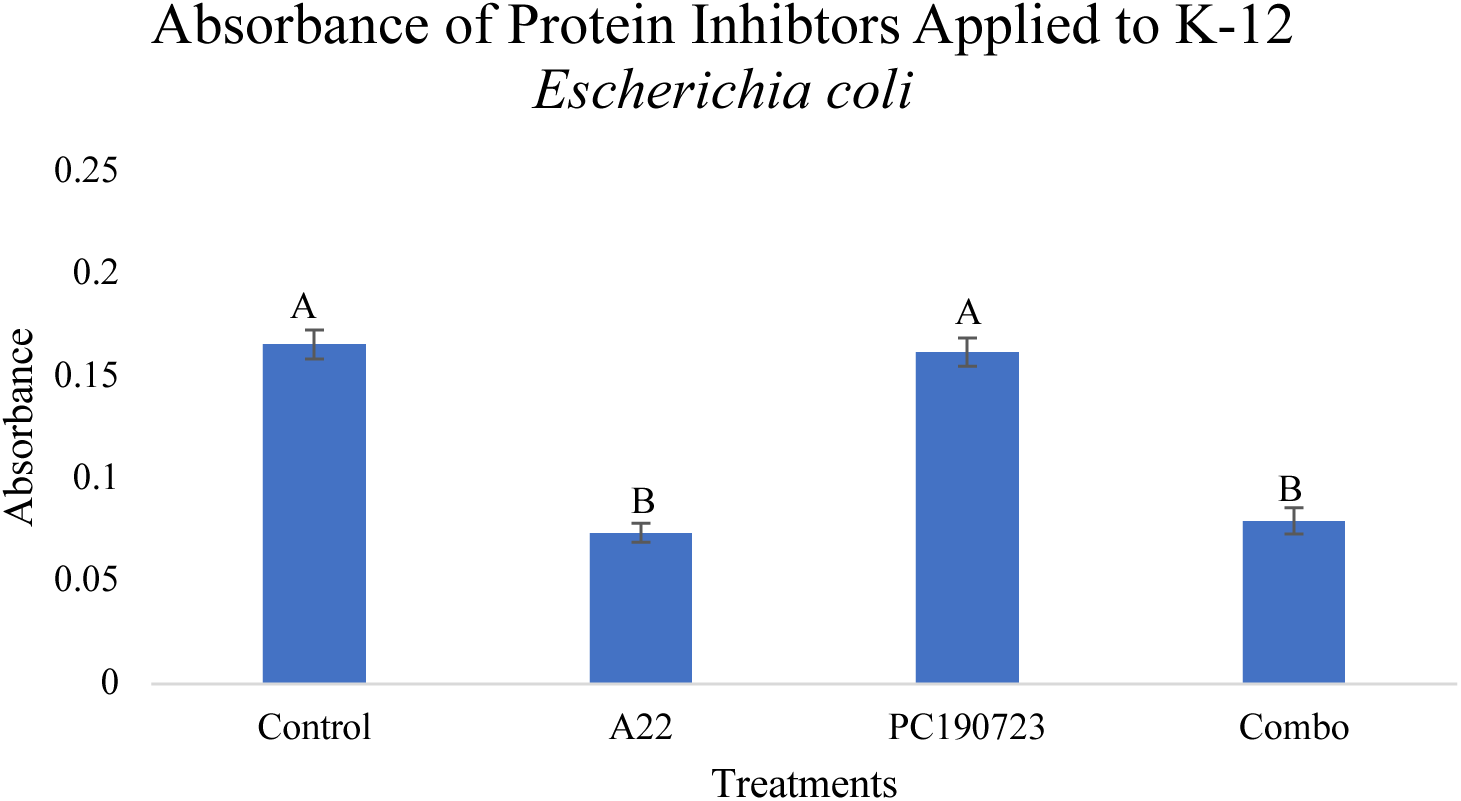
Mean absorbances of various antimicrobial treated well plates. The treatments are the following: Control, A22 (200 μg/mL, PC190723 (100 μg/mL), and Combination (A22: 200 μg/mL, PC190723: 100 μg/mL). Error bars represent one standard error from the mean. Letters represent Tukey’s mean separation.

### Antibiotic Experiment

After testing the antibiotic susceptibility of A22-treated *E. coli*, it was found that A22 does not increase nor decrease the antibiotic susceptibility. The effect of A22 treatment was indistinguishable from the Amoxicillin treatment. Similarly, a combination of the two treatments was not significantly different from either treatment, individually. All three treatments differed from the control. See Figure 9 and Table 4.

**Table 4.** ANOVA results from Antibiotic Experiment. Asterix (*) represent significant values (p<.05).

| *Error* | *df* | *SS* | *MS* | *F* | *P* |
| --- | --- | --- | --- | --- | --- |
| Model | 3 | 0.07486778 | 0.024956 | 4.6169 | <.0053* |
| Treatment | 3 | 0.0032582 |  | 3.0315 | <.0352* |
| Error | 68 | 0.36756400 | 0.005405 |  |  |
| Total | 71 | 0.44243178 |  |  |  |

**Figure 9.**
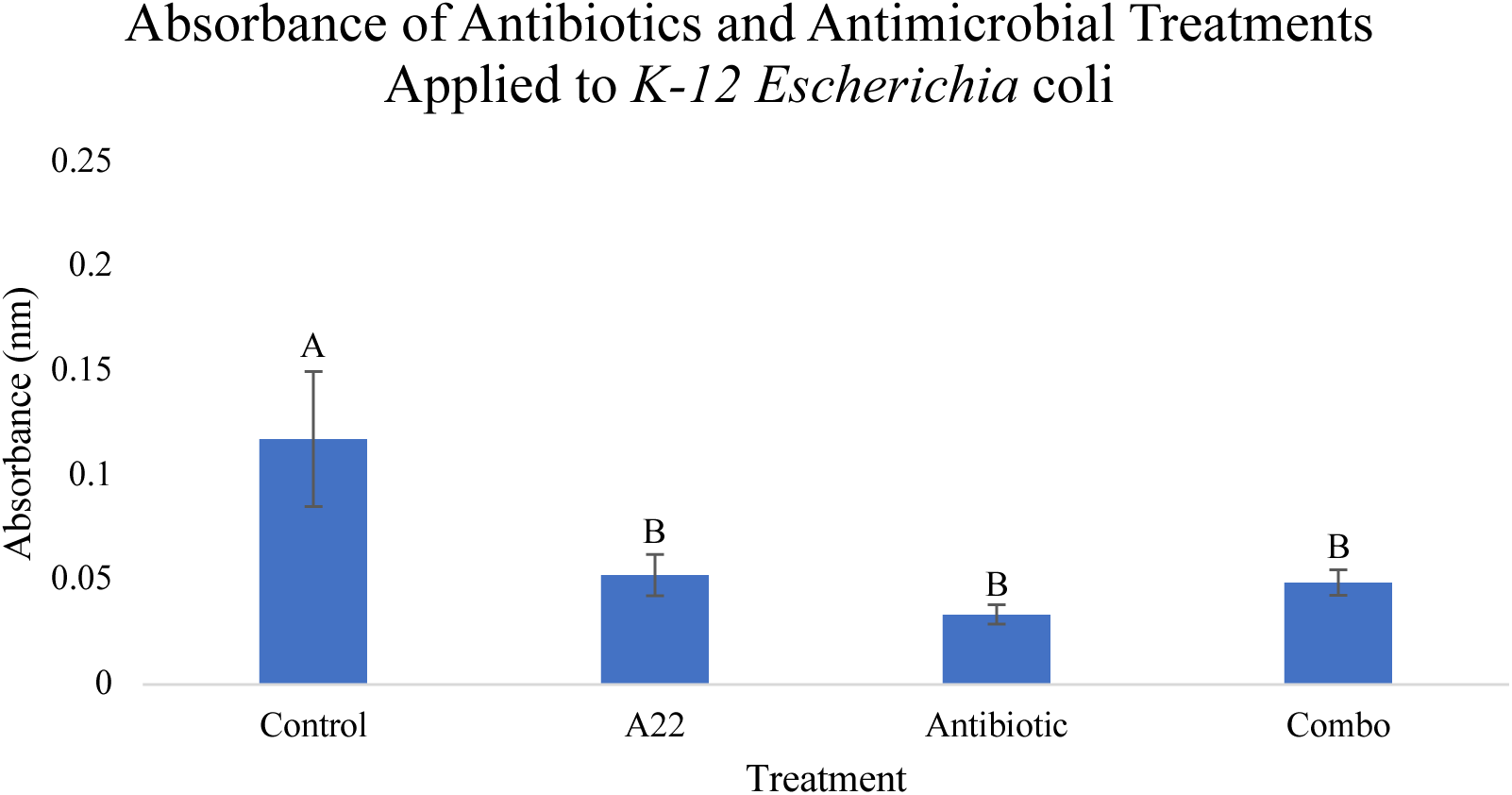
Mean absorbances of various antimicrobial treated well plates. The treatments are the following: Control (Zero), A22 (200 μg/mL), Antibiotic (2 μg/mL), and Combination (A22: 200 μg/mL, Antibiotic: 2 μg/mL). Error bars represent one standard error from the mean. Letters represent Tukey’s mean separation.

### Microscopy

From qualitative analysis of pictures taken, it was shown that A22 did, in fact, change cells transform cells from a rod-like structure to a more circular one. Cells treated with PC190723 showed no detectable changed under a light microscope. See Figures 10A and 10B.

**Figure 10.**
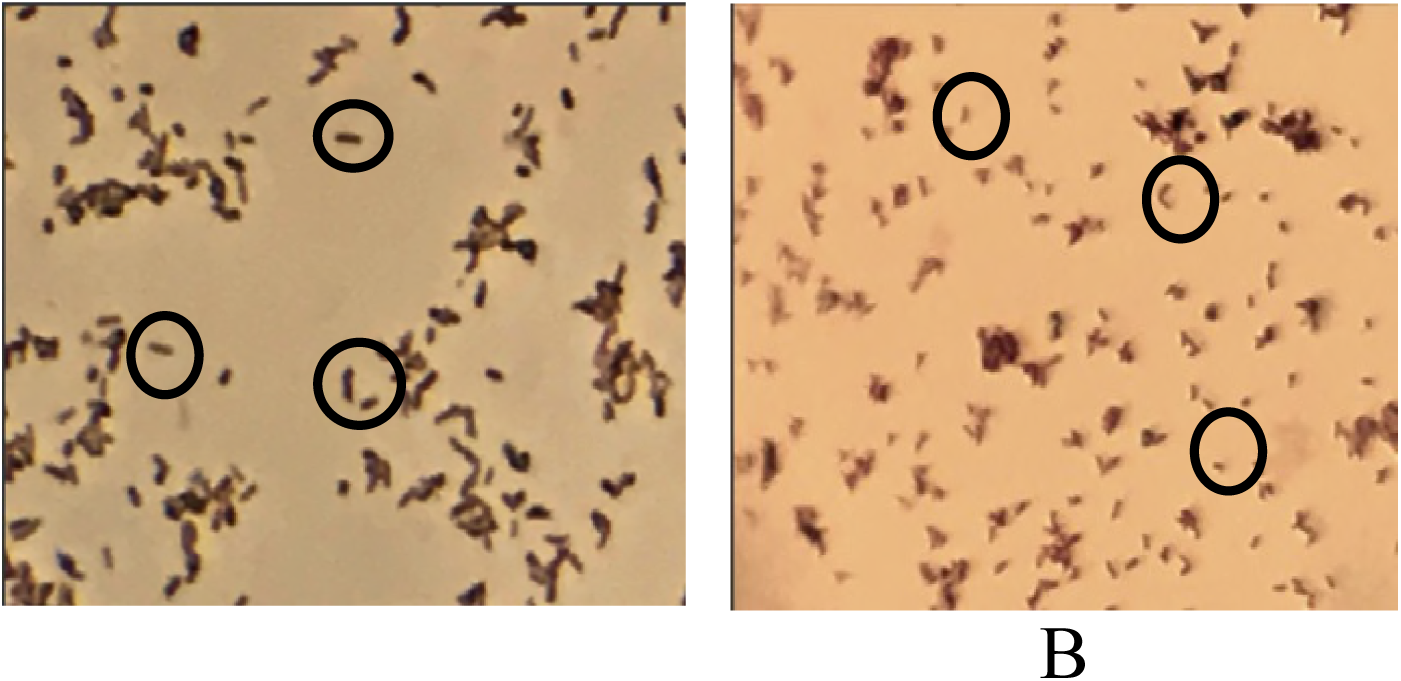
A: *E. coli* cells prior to A22 treatment; cells are seen to be rods. B: *E. coli* cells after A22 treatment; cells are seen to be more circular, bent, contorted, and smaller. Picture is taken at 400x

## Discussion

The results of this experiment confirmed that A22 was significantly better than the control in its ability to inhibit biofilm formation. There was no significant effect of PC190723 relative to the control and no dose response to PC190723. When analyzing the modes of action of these two protein inhibitors, it is rational for A22 to be more effective. A22 inhibits MreB within the bacteria which directly controls for the rod shape of cells, similar to the role of the protein actin in eukaryotes. When MreB is no longer active, cells became circular which indirectly affected the establishment of the biofilms on the surface. This conclusion cannot be made without examining the role of PC190723. Within the cell, ftsZ acts to regulate the plane of cell division, similarly to the protein tubulin in eukaryotic cells. FtsZ certainly does play a role in colony development, as it initially localizes the site of bacterial cell division (Margolin, 2005), but this role may not significantly affect the biofilm density. The inhibition of ftsZ by PC190723 could potentially disrupt the cell division process, but due to the complexity of this process, it cannot be the only protein responsible for regulation. Other important proteins involved in this process may be able to compensate for the inhibition of PC190723 and simply continue with the process as normal (Rhind & Russell, 2012). Because there was no significant reduction in biofilm density when using PC190723, greater concentrations of the chemical could be tested to see if this lack of response may simply be due to weak concentrations of treatment.

When examining the mid-level dosage of A22, which significantly decreased biofilm density, it is plausible for further studies to be done to find the exact MIC. What is determined from this experiment is that the MIC lay somewhere between the low dosage (100 µg/mL) and the medium dosage (200 µg/mL).

The microscopy experiment confirmed that A22 was having the expected effect on the cell shape. As it should, the chemical was properly binding to the MreB binding site and not allowing cells to polymerize the naturally occurring MreB. This molecular change is what shifts the cells from a bacillus shape to a coccoid one.

The second experiment holds a predicable result based on the result of the first experiment. It is reasonable to think that there would be no synergistic effect of the two chemicals when one of them, PC190723, has no effects on its own. The results from this experiment further confirm that PC190723 did nothing to affect biofilms (at least in the concentration used). The treatment had no observable effect when applied individually and continued this trend when applied in combination with A22. This result was expected after examining the data from the first experiment, yet it was contrary to the original hypothesis which proposed that the effect on biofilms would be the least in the PC190723 treatment, at a middle level in the A22 treatment, and the greatest in the combination experiment.

This combination experiment was repeated twice due to the question of the A22 treatment masking any additive effect of PC190723 that may emerge. When the same results were obtained during the second trial, the results of the first trial were accepted. When the treatments are combined, the cells, in theory, transform into a circular shape and have a disrupted process of division. Specifically, the plane of division will be altered and shifted to a different plane. If the cells are truly spherical, then a disrupted plane of division will not have an effect as all division lines will equally splice the cell. This may be one reason why no synergistic effect emerged within the Combination experiment. A further study down this line of experimentation would be to use a stronger dosage of PC190723 that has been pre-determined to have an effect.

The final experiment in the sequence of this study is the antibiotic experiment. This experiment allowed insight into the true question of the study; can biofilms be inhibited well enough to allow for antibiotic susceptibility to increase? The data for this experiment led to the conclusion that A22 did not have an effect on antibiotic susceptibility. While the three treatments differed significantly from the control, the treatments were indistinguishable from each other. These results may in part be due to a general lack of growth in the *E. coli* culture, with the control Absorbance being noticeably lower than previous experiments. Additionally, a lower dosage of Amoxicillin could be tested in future experiments as 2 µg/mL may have simply killed all biofilms rather than inhibiting their initial formation. A lower dosage of antibiotics would allow for less general damage done by broad-spectrum antibiotics while also allowing for a potential effect on biofilm degradation. Although, despite the hypothesis being disproved, the experiment does allow insight on an inhibitory compound which is as equally effective as a widely used antibiotic.

With these conclusions in mind, a potential mechanism of biofilm attachment can be proposed. When biofilms form, they form in layered, dense mats. This layering is easily done by cells of the proper shape, such as the rod-like shape of *E. coli* biofilm cells. The layered cells can 1) easily attach to each other and 2) easily attach to the site of infections, providing a strong base of strength and attachment throughout the film. Though, when a morphological change is induced, such as the one caused by A22, cells may no longer inter-attach and attach to the site of infection in the way they naturally do. While, this does not influence antibiotic susceptibility, as shown in the Antibiotic experiment, it may influence other factors within the treatment of bacterial infections. Future investigations looking into this mechanism of strength and attachment could provide conclusive results in the deterioration of biofilms and their weaknesses.

## Acknowledgements

I would like to first thank Dr. Sheck for advising me and guiding me through the research process. Thanks to her, I have been able to not only find my scientific passions but also pique my inquisitive nature and problem-solving abilities. Thank you to Dr. Monahan for advising me throughout the summer data collection and for the continuous feedback she has given me. Thank you to the NCSSM Research in Biology class of 2019 for helping me along my research path, and to the NCSSM Research in Biology class of 2018 for putting forth the time to mentor me.

Thank you to Kevin Zhang and Tyler Edwards for being wonderful lab assistants over the summer, aiding me in countless ways. Finally, I would like to thank the North Carolina School of Science and Mathematics and the GlaxoSmithKline Endowment to NCSSM for allowing me the amazing opportunity to conduct research for the first time. It has been an incredible experience and I have gained skills that I am confident I will carry with me for the remainder of my research career as well as beyond.

## Literature Cited

Anderl, J. N., M. J. Franklin, and P. S. Stewart. 2000. Role of antibiotic penetration limitation in Klebsiella pneumoniae biofilm resistance to ampicillin and ciprofloxacin. Antimicrobial Agents and Chemotherapy 44: 1818–1824. https://doi.org/10.1128/AAC.44.7.1818-1824.2000

Andreu, M., C. Schaffner-barbero, S. Huecas, D. Alonso, M. L. Lopez-rodriguez, L. B. Ruiz-avila, and R. Nu. 2010. The Antibacterial Cell Division Inhibitor PC190723 Is an FtsZ Polymer-stabilizing Agent That Induces Filament Assembly. The Journal of Biological Chemistry 285: 14239–14246. https://doi.org/10.1074/jbc.M109.094722

Barsoumian, A. E., K. Mende, C. J. Sanchez, M. L. Beckius, J. C. Wenke, C. K. Murray, and K. S. Akers. 2015. Clinical infectious outcomes associated with biofilm-related bacterial infections: A retrospective chart review. BMC Infectious Diseases 15: 1–7. https://doi.org/10.1186/s12879-015-0972-2

Bonez, P. C., G. G. Rossi, J. R. Bandeira, A. P. Ramos, C. R. Mizdal, V. A. Agertt, … M. M. A. de Campos. 2017. Anti-biofilm activity of A22 ((S-3,4-dichlorobenzyl) isothiourea hydrochloride) against Pseudomonas aeruginosa: Influence on biofilm formation, motility and bioadhesion. Microbial Pathogenesis 111: 6–13. https://doi.org/10.1016/j.micpath.2017.08.008

Brown, D. R., P. J. Baker, V. V Barynin, D. W. Rice, and S. E. Sedelnikova. 2008. Selective Anti-Staphylococcal Activity. Science Volume 321: 1673–1676.

CDC. 2013. Antibiotic resistance threats in the United States, 2013. Current 114. https://doi.org/CS239559-B

Costerton, J. W. Philip S. Stewart, E. P. G. 1999. Bacterial Biofilms: A Common Cause of Persistent Infections. Science 284: 1318–1322. https://doi.org/10.1126/science.284.5418.1318

D. Zeng, Personal Communication, March 26, 2018

Hoyle, B. D., J. Alcantara, and J. W. Costerton. 1992. Pseudomonas aeruginosa biofilm as a diffusion barrier to piperacillin. Antimicrobial Agents and Chemotherapy 36: 2054–2056. https://doi.org/10.1128/AAC.36.9.2054

IDEXX Reference Laboratories. 2013. Microbiology Guide to Interpreting Minimum Inhibitory Concentration (MIC) 1-4. Retrieved from http://www.idexx.co.uk/pdf/en_gb/smallanimal/reference-laboratories/testmenu/GuidetoInterpretingMICs.pdf

Kumar, A., A. Alam, M. Rani, N. Z. Ehtesham, and S. E. Hasnain. 2017. Biofilms: Survival and defense strategy for pathogens. International Journal of Medical Microbiology 307: 481–489. https://doi.org/10.1016/j.ijmm.2017.09.016

Margolin, W. 2005. FtsZ and the division of prokaryotic cells and organelles. Nature Reviews Molecular Cell Biology 6: 862. Retrieved from http://dx.doi.org/10.1038/nrm1745

Rasamiravaka, T., Q. Labtani, P. Duez, and M. El Jaziri. 2015. The formation of biofilms by pseudomonas aeruginosa: A review of the natural and synthetic compounds interfering with control mechanisms. BioMed Research International 2015. https://doi.org/10.1155/2015/759348

Rhind, N., and P. Russell. 2012. Signaling pathways that regulate cell division. Cold Spring Harbor Perspectives in Biology 4: 1–15. https://doi.org/10.1101/cshperspect.a005942

